# Antimicrobial Resistance in Diverse Urban Microbiomes: Uncovering Patterns and Predictive Markers

**DOI:** 10.1101/2024.03.08.584116

**Authors:** Rodolfo Brizola Toscan, Wojciech Lesiński, Piotr Stomma, Balakrishnan Subramanian, Paweł P. Łabaj, Witold R. Rudnicki

## Abstract

Antimicrobial resistance (AMR) poses a significant global health threat, exacerbated by urbanization and anthropogenic activities. This study investigates the distribution and dynamics of AMR within urban microbiomes from six major U.S. cities using metagenomic data provided by the CAMDA 2023 challenge. We employed a range of analytical tools to investigate sample resistome, virome, and mobile genetic elements (MGEs) across these urban environments. Our results demonstrate that AMR++ and Bowtie outperform other tools in detecting diverse and abundant AMR genes, with binarization of data enhancing classification performance. The analysis revealed that a portion of resistome markers is closely associated with MGEs, and their removal drastically impacts the resistome profile and the accuracy of resistome modeling. These findings highlight the importance of preserving key MGEs in resistome studies to maintain the integrity and predictive power of AMR profiling models. This study underscores the heterogeneous nature of AMR in urban settings and the critical role of MGEs, providing valuable insights for future research and public health strategies.

## 1 INTRODUCTION

Antimicrobial resistance (AMR) is a phenomenon that arises when bacteria, viruses, fungi, and parasites undergo genetic changes, rendering them insensitive to the effects of antimicrobial agents, thereby making infections more difficult to treat and increasing the risk of disease transmission, morbidity, and mortality (Sun et al. (2022)). The emergence and spread of AMR are inherently driven by anthropogenic factors, and it is estimated that over one million people died due to AMR in 2019 (Christopher J L Murray (2022)). In conjunction with suboptimal wastewater treatment processes that fail to degrade residual antibiotic agents, inappropriate use of antibiotics has led to a persistent increase in the environmental abundance of antimicrobial compounds (Aden and Bashiru (2022)).

Antimicrobial resistance genes (ARGs) are transmitted either vertically through binary fission in bacteria or horizontally through horizontal gene transfer (HGT) mechanisms, including conjugation, transformation, and transduction (Ochman et al. (2000)). Transformation involves bacterial uptake of genetic material from their surroundings, while conjugation involves the direct exchange of genetic material between bacterial cells. Unlike these processes, transduction is mediated by viruses and mobile genetic elements, highlighting the necessity of considering these dynamic entities when investigating antimicrobial resistance (Canchaya et al. (2003); Frost et al. (2005)).

Resistome profiling, particularly in hotspots like wastewater treatment facilities, meat processing plants, hospitals, and urban areas, has gained significant attention (Karkman et al. (2018); Honda et al. (2023); Aarestrup (2012); Mulvey and Simor (2009); Vassallo et al. (2022)). Human activities impact the resistome substantially (Vassallo et al. (2022)). The global scientific community has thus intensified efforts to understand resistome dynamics (Cobo-DÍaz et al. (2021)). One notable initiative is MetaSUB, which periodically sequences metagenomic material from urban public spaces such as metro stations and bus stops (Ryon et al. (2022)). Metagenomics, advantageous over culturing methods for its independence from known genes and primers, relies on comprehensive reference databases like CARD (Alcock et al. (2022)) and Resfam (Gibson et al. (2014)) for ARG identification.

This study extensively analyses 143 urban environmental metagenomic samples (see Table 1) and antibiotic susceptibility data from 145 hospital patients and their delivered isolates.

**Table 1.** Overview of Sample Distribution by City for the Study. The table lists each city with its corresponding ID and the number of samples collected.

| ID | City | Sample number |
| --- | --- | --- |
| BAL | Baltimore | 14 |
| DEN | Denver | 45 |
| MIN | Minneapolis | 6 |
| NYC | New York | 46 |
| SAC | Sacramento | 16 |
| SAN | San Antonio | 16 |

Both datasets, i.e., the metagenomic fastq files and the isolates’ resistome profiles, were provided by the CAMDA 2023 Challenge (2023) organization team. The metagenomic samples are an arbitrary subset of the MetaSUB sequencing pool, where initial analysis has led to drafting a global metagenomic map of urban microbiomes and antimicrobial resistance (The International MetaSUB Consortium (2021)). These samples were collected from six major U.S. cities (Danko et al. (2021)). Our investigation encompasses their resistome, virome, and mobile genetic elements, employing a diverse array of techniques both independently and in conjunction (see Figure 1).

**Figure 1.**
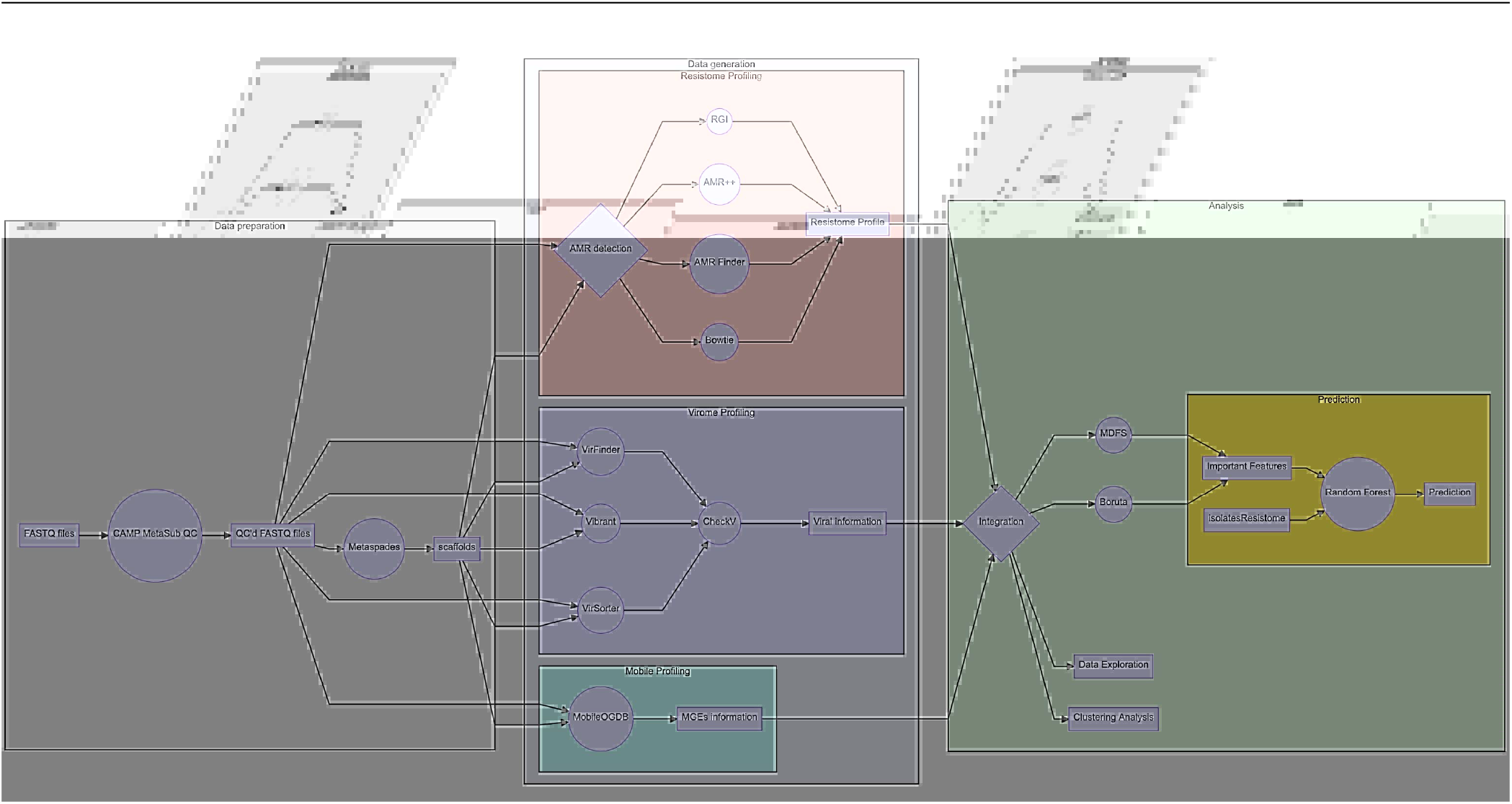
This diagram provides a high-level view of the methods workflow used in this study, illustrating the stages from data preparation through to prediction. The process includes quality control of sequencing data, various profiling methods for resistome, virome, and mobile genetic elements, followed by integration and analysis using feature selection and machine learning techniques.

We employed mathematical modelling and statistical techniques to analyze the metagenomic data and predict the origins of the samples. We utilized random forest classifiers for feature selection and classification, leveraging algorithms like Boruta and Multi-Dimensional Feature Selector (MDFS) to identify key resistome markers. Additionally, we computed cosine similarities between samples based on their antimicrobial resistance (AMR) profiles and applied clustering algorithms to explore the data structure. Singular Value Decomposition (SVD) was used for dimensionality reduction to enhance the accuracy of similarity calculations (Berry et al. (1995)). Furthermore, we investigated the association between mobile genetic elements (MGEs) and AMRs, conducting filtering experiments to assess the impact of MGE-associated AMRs on resistome profiles.

These mathematical approaches allowed us to derive meaningful insights into the distribution and dynamics of AMR in urban microbiomes. By mapping resistome profiles across different urban environments and evaluating the precision and applicability of various resistome profiling tools, our study significantly contributed to the advancement of resistome analysis methodologies. Despite facing challenges, our findings underscore the critical role of MGEs in resistome studies and provide valuable insights for future research and public health strategies.

## 2 METHODS

### 2.1 **Data Preparation**

The study utilized fastq files obtained from a publicly available repository of MetaSUB data (www.metasub.org) accessible through CAMDA 2023 Challenge (2023) page. A total of 143 libraries from six different cities in the United States were examined (Table 1). To ensure the dataset’s quality, the MetaSUB-CAMP metagenomic tool suite Tierney et al. (2023) was employed. This suite conducted quality control procedures, including the removal of host reads (Genome Reference Consortium Human Build 38 - RefSeq assembly accession: GCF 000001405.26) and low-quality sequences. The quality-controlled reads were then subjected to *de novo* assembly using metaSPADES Nurk et al. (2017) with standard parameters.

### 2.2 **Data Generation**

#### 2.2.1 Resistome profiling

We constructed resistome profiles using four different analytical methods. For short quality-controlled reads, we used AMR++ v3.0 (Microbial-Ecology-Group (2023)) and Bowtie v2.5.1 (Langmead and Salzberg (2012)). AMR++ makes use of MEGARes v3.0, a comprehensive AMR database with an acyclic hierarchical annotation structure (Bonin et al. (2022)). For the Bowtie approach, we aligned the reads against a custom database with Bowtie2 using standard parameters. This database contained a comprehensive collection of indexed antimicrobial resistance genes, combining sequences from the Comprehensive Antibiotic Resistance Database (CARD) (Alcock et al. (2022)) with an additional set of manually curated genes, kindly provided by Dr. Nelly Selém (Mojica (2023)).

For elongated, assembled reads, we utilized two other methods: AMRFinderPlus v3.11.14 (Feldgarden et al. (2021)) and Resistance Gene Identifier (RGI) V3.3.1, developed by and using the CARD initiative (Alcock et al. (2022)). In executing AMR++, AMRFinderPlus, and RGI, we adhered to the standard pipelines without any modifications to the databases or pipeline parameters.

#### 2.2.2 Resistome normalization

Considering the contrasting sequencing depth across the dataset, we employed a range of normalization techniques for gene counts. These counts were normalized against the following parameters: 1) the quantity of quality-controlled base pairs, 2) the total number of assembled base pairs, 3) the detected SSU count, and 4) the count of SSUs exhibiting a minimum of 50% coverage.

Our examination of assembled contigs for Small Subunit (SSU) identification involved the utilization of the bbduk.sh tool from the BBTools suite (Bushnell (2021)). We referenced the SILVA Small Subunit database (release 138.1) for this purpose Quast et al. (2013). The detection of SSU fragments was quantified at varying coverage thresholds, including any detectable level (above 0%) and a more stringent criterion of over 50% coverage.

#### 2.2.3 MGE identification, annotation and classification

We used MobileGo-DB (Brown et al. (2022)) for the MGE identification. MobileGo-DB serves as a meticulously curated database housing well-documented protein sequences of Mobile Genetic Elements (MGEs), encompassing diverse elements such as transposons, plasmids, integrons, and various other mobile entities. MGE identification using mobileGo-DB comprises of three steps:

1. Identification of open reading frames using Prodigal (Hyatt et al. (2010)).
2. Creation of alignment summaries to a mobile orthologous groups database file utilizing Diamond (Buchfink et al. (2015)).
3. Compilation of element-mapping data, providing a summary of matches to proteins from various mobile element classes.

In the initial step, we identified open reading frames within the assembled contigs sourced from US cities. This identification process used the Prodigal tool, as outlined in the study conducted by Hyatt et al. (2010).

Progressing to the second stage, we generated summaries illustrating the alignments between the genetic material and a specialized mobile orthologous groups database. The efficiency of this alignment was ensured through the use of the Diamond tool, as elucidated in the research by Buchfink et al. (2015). Finally, in the third phase, we process the results, offering a summary of matches discovered in the samples to proteins from different classes of mobile genetic elements.

#### 2.2.4 MGE quantification and AMR association

We identified mobile genetic elements (MGEs) and computed the frequency of MGE genes across the dataset using the MGE gene hits. Our analysis focused on MGE genes found in more than 50% of the samples from each city. To investigate the association between MGEs and antimicrobial resistance (AMR), we revisited the original genetic data, searching for AMR genes proximal to the MGE genes. MGEs and resistome markers co-located on the same contig were considered associated and were categorized as mobile antimicrobial resistance markers (mAMRs).

#### 2.2.5 Virome profiling

Virome profiling was conducted using three distinct tools, each chosen for its specific utility in viral genome identification and analysis. VirSorter v2.2.4 (Guo et al. (2021)), a widely used tool, employs prophage sequences to identify virus-like signatures in microbial datasets. VirFinder (Ren et al. (2017)) v1.1 is known for its statistical learning approach, which assigns a likelihood score to sequences for their viral origin, enhancing detection specificity. Vibrant v1.2.1 (Kieft et al. (2020)) utilizes machine learning and known viral databases to annotate and predict viral sequences with high accuracy. The outputs from VirSorter, VirFinder, and Vibrant were combined, and duplicate entries were removed to retain only unique viral sequences. Subsequently, the unique sequences underwent a quality control process using CheckV (Nayfach et al. (2021)), which assesses the completeness and quality of the detected viral genomes, ensuring that the data used in further analyses are of high integrity. These tools were integrated into the Snakemake viral investigation pipeline (MetaSUB-CAMP (2024)), which was operated with standard parameters to ensure consistent and reproducible analysis across datasets.

#### 2.2.6 K-mer Profiling

K-mer profiling was conducted to assess the diversity and complexity of the metagenomic samples. We used Jellyfish (Marcais and Kingsford (2011)) to compute k-mer statistics, generating a 143x12 table, with one row per sample and four columns for each k-mer size (33, 55, and 77). The k-mer sizes of 33, 55, and 77 were chosen to balance sensitivity and specificity in sequence detection. Shorter k-mers (e.g., 33) detect a broader range of sequences and capture small genetic variations, while longer k-mers (e.g., 77) offer higher specificity and reduce false positives by ensuring unique matches. This multi-scale approach leverages the benefits of different k-mer lengths, as supported by previous genomic analysis studies (Chikhi and Medvedev (2014); Li and Wang (2010)).

The metrics calculated included unique, distinct, total, and max count for each k-mer size. The *unique* metric counts k-mers occurring exactly once, serving as a direct indicator of sample diversity. The *distinct* metric counts the number of k-mers while ignoring their multiplicity, representing the cardinality of the set of k-mers. The *total* metric sums the occurrences of all k-mers, reflecting the overall abundance and richness of the sample. The *max count* metric identifies the highest occurrence of any single k-mer within a sample, indicating the presence of highly repetitive sequences or dominant species.

#### 2.2.7 Data Integration

All intermediate analysis results from the profiling of resistome, mobilome, virome, and k-mer counts were processed and wrangled for subsequent interpretation and analysis through clustering, modeling, and prediction techniques. The results generated by each tool was compiled into matrices and analysed using the R statistical programming language.

### 2.3 **Analysis**

#### 2.3.1 Analysis of AMR-based city similarity

The initial goal was to perform an exploratory analysis of the clustering structure of the urban samples derived from their AMR profiles. To this end, clustering-based approaches were tested, based on *L*2 norm cosine similarity matrix (Horn and Johnson (1985)) computed using absence-presence tables of AMRs. It is expected that if AMR levels are related to the geographical location of the samples, then by using AMR-based similarity to cluster the samples, one could recover each sample’s city assignments by inspecting the cluster labels. However, after initial tests, we discovered that the clustering structure cannot be easily mapped to the original city labels. Therefore a more fundamental approach was used, that was concerned with the AMR-based sample similarities themselves, not the clusters built upon them.

The cosine similarity between samples was computed for each AMR profiling approach: AMRFinderPlus, AMR++, RGI and Bowtie. Then, a statistical analysis of the relationship between the values of the similarities and city labels was performed. Several variants of the cosine-based similarities were computed and compared. Differences between the similarities inside vs across cities were examined, and their statistical significance was assessed. Our protocol was aimed to answer the question of ”Does the AMR- based sample similarity carry information about the geographical origin of the samples?”. The general outline of the protocol is stated below:

1. For each tool, compute the sample similarities.
2. For each tool, based on sample similarities, compute summary statistics comparing the inside-city similarity of the samples with the between-city similarity.
3. Assess the statistical significance of the differences and compare the statistics across different AMR finding tools and similarity variants.

#### 2.3.2 Cosine similarity calculation

For each tool, different variants of the cosine similarities were computed. Each variant tested starts with a raw absence/presence table. It is then used to compute plain cosine similarity. Optionally, a combination of the following transformations was applied to the data between those two elementary steps. Details of each step are discussed in the next parts of the manuscript:

*•* **Input markers filtering**: use all available markers or only those relevant for the decision variable ‘city‘.
*•* **Usage of SVD embedding**: either apply SVD embedding (Wall et al. (2003); Berry et al. (1995)) on the absence/presence table or don’t.
*•* **Sparsification of the cosine similarity matrix**: either zero out weak connections or don’t.

Transformations were applied in the order they are listed. Effects of each combination of these transformations were tested.

#### 2.3.3 Transformation details

For selecting the markers related to the city indicator, relevance was computed by a chi-squared test based on mutual information (*MI*) between the decision and the features (Mnich and Rudnicki (2020)). We have used a custom function mimicking the behaviour of the MDFS 1-D feature selector (Piliszek et al. (2019)) (generalized for handling non-binary decision variables, that is, the ’city indicator’). For the SVD embedding, we have used the *UD* part of the *SV D* decomposition (Berry et al. (1995)) applied to the marker matrix *M*, where *M* = *UDV*, where rows of *M* correspond to samples, and columns to the markers. Each row of *UD* matrix represents the joint information of the each sample contained in its AMR levels, while first *k* columns carry the most variation across different samples.

Sparsification step zeroes out weak connections, according to the weight threshold chosen by clique counting on the thresholded graph. A clique is a subset of vertices in a graph such that every two distinct vertices are connected by an edge. We choose threshold for which the number of observed cliques of size at least 3 is maximal. Such threshold is dependent on the structure of the graph, thus it varies between variants. This is a heuristic we found empirically work well in various scenarios. More details can be found in the supplementary material.

#### 2.3.4 Summary statistics of the sample similarities

Here we describe a simple statistic used to assess how, on average, well separated are samples coming from different cities. We have computed the similarity for each of <u>^1^</u> *N* (*N −*1) pairs created from *N* samples. Set of computed similarities can be partitioned into the set of ’inside city’ similarities *S_in_* and ’between city’ similarities *S_btw_*. Similarities between samples were computed using levels of AMRs. If those levels are overall relevant for the geographical location of the samples, we would expect, on average, for the ’inside city’ similarity to be bigger than the ’between city’ similarity. Therefore, we found it meaningful to compute the following summary statistic that summarizes each similarity matrix:

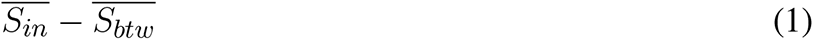

where *A* denotes mean of elements in set *A*. The greater the value, the greater the similarities between samples from the same city than the similarities between samples from different cities.

### 2.3.5 Statistical significance assessment

To adjust for possible randomness of the differences between the computed summary statistics, we have utilized both common non-parametric approaches: resampling-based point estimate with uncertainty estimation, as well as permutation-based significance test where applicable (Efron and Tibshirani (1993)).

To estimate the standard error of the statistic, we have used a variation of leave-*d*-out jackknife (Shao and Wu (1989)). In standard leave-*d*-out jackknife, one computes the statistic for all (or random sample of all) possible subsets of samples of the original dataset (size *N*) that have *N − d* elements, and uses the resulting replicates to compute the spread of original statistic. In our case, we used a stratified variant of such procedure because of a serious class imbalance in the variable ’city’. We want to leave out <u>^1^</u> samples in each subset. Therefore, to ensure equal treatment of each class of the ’city’ variable, to compute each replicate, we leave 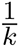 out of each class.

We have also used a permutation test (Efron and Tibshirani (1993)) for the statistic 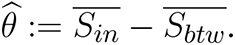.

More detailed information on specific parameters used in the procedures desribed above can be found in the supplementary material.

#### 2.3.6 Feature selection and classification

We used two methods for feature selection: Boruta algorithm (Kursa and Rudnicki (2010)) and Multi Dimensional Feature Selector (MDFS) (Mnich and Rudnicki (2020); Piliszek et al. (2019)). The feature selection and random forest classification focused on predicting the origin of the samples, specifically the ‘city’ indicator. Boruta uses the importance score from multiple runs of the RF algorithm to discover the informative variables. In each iteration, the original data set is extended by adding a randomized copy of each variable, MDFS is based on information theory and considers synergistic interactions between descriptive variables. The algorithm returns binary decisions about variables’ relevance and ranking based on Information Gain and p-value.

For modelling, we used **Random Forest** algorithm Breiman (2001), which is based on decision trees and works well *out of the box* on most data sets Fernández-Delgado et al. (2014). It also obtains relatively well results with a small number of observations. The classifier was used in three ways: all versus all, one versus all, and one versus one. Models were evaluated by accuracy (multiclass cases) and by Matthews Correlation Coefficient (MCC) Matthews (1975) and area under the receiver operating curve (AUROC or AUC) for binary classification.

## 3 RESULTS

### 3.1 **Data description and interpretation**

#### 3.1.1 On the resistome

Table 2 provides a summary of AMR detection counts using four different methods: AMR++, Bowtie, RGI, and AMRFinder. AMR++ identified the highest total count of unique genes at 977, with an average (mean) of 68 and a standard deviation of 94, indicating considerable variability in the data. Bowtie detected a total of 342 unique genes, with a mean of 67 and a standard deviation of 57 – indicating more moderate spread around its mean. RGI identified 252 unique genes, with a lower mean of 12 and a standard deviation of 17, reflecting less variability compared to AMR++ and Bowtie. AMRFinder, while detecting the fewest unique genes at 142, had a mean of 9 and a standard deviation of 12, indicating a relatively consistent detection rate.

**Table 2.** Comparison of AMR Detection Methods. This table summarizes the total counts, mean values, standard deviations, and variability of antibiotic resistance detection across four methods used to AMRs in this study. Variability was calculated as Standard Deviation divided by Mean.

| Method | Total | Mean | Standard Deviation | Variability |
| --- | --- | --- | --- | --- |
| AMR++ | 977 | 67.8 | 93.5 | 1.4 |
| Bowtie | 342 | 67.2 | 57.1 | 0.9 |
| RGI | 252 | 12.0 | 17.1 | 1.4 |
| AMRFinder | 142 | 9.4 | 11.4 | 1.2 |

Overall, AMR++ and Bowtie both show higher mean and standard deviation counts of detected genes when compared to RGI and AMRFinder. AMR++ and RGI both display similar level of variability, as computed by dividing standard deviation by mean values.

We performed a correlation analysis between the number of genes found per sample for each of the tools and the number sequenced nucleotides and assembled basepairs. None of the methods displayed a significant correlation (*>*55%), suggesting that the detection of AMR markers is largely independent of sample size.

The bar plot on Figure 3 shows the total number of normalized genes per class across different cities, with gene classes represented by Biocides (coral), Drugs (dark green), Metals (cyan), and Multi-compound (purple). The cities included are Baltimore (BAL), Denver (DEN), Minneapolis (MIN), New York City (NYC), Sacramento (SAC), and San Francisco (SAN). Notably, New York City (NYC) exhibits a significantly higher count of normalized genes in the Drugs class compared to other cities, indicating a prominent presence of drug-related genes. Denver (DEN) and Sacramento (SAC) also display noticeable counts in the Drugs class, though much lower than NYC. The other cities (BAL, MIN, SAN) have relatively lower and more balanced counts across different gene classes. The presence of Biocides, Metals, and Multi-compound genes is generally low across all cities. This figure underscores the variability in the distribution of gene classes among different urban environments, with a notable emphasis on the predominance of drug-related genes in New York City.

**Figure 2.**
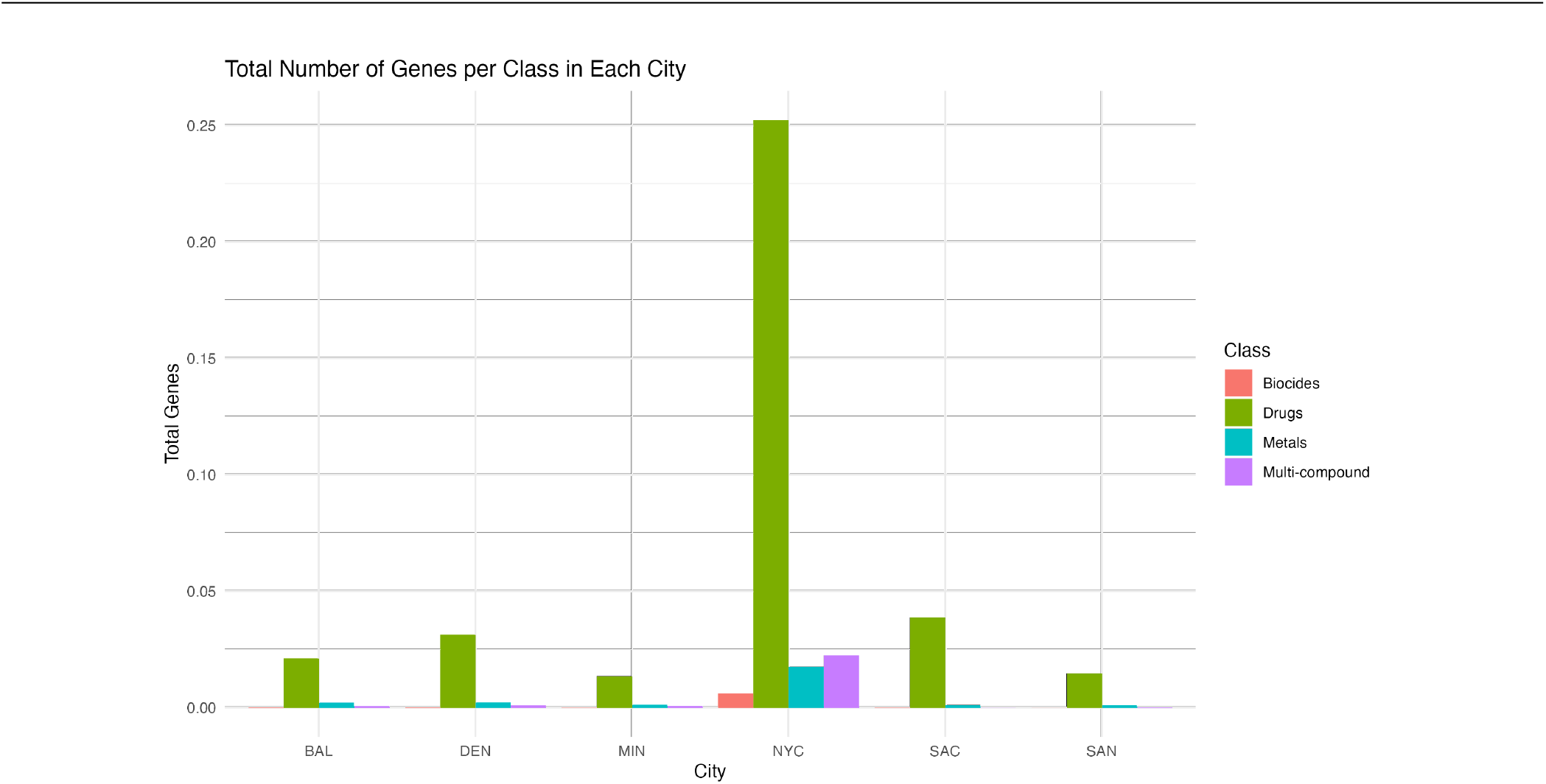
The bar plot shows the total number of normalized resistome marker detected by AMR++ per class across different cities.

**Figure 3.**
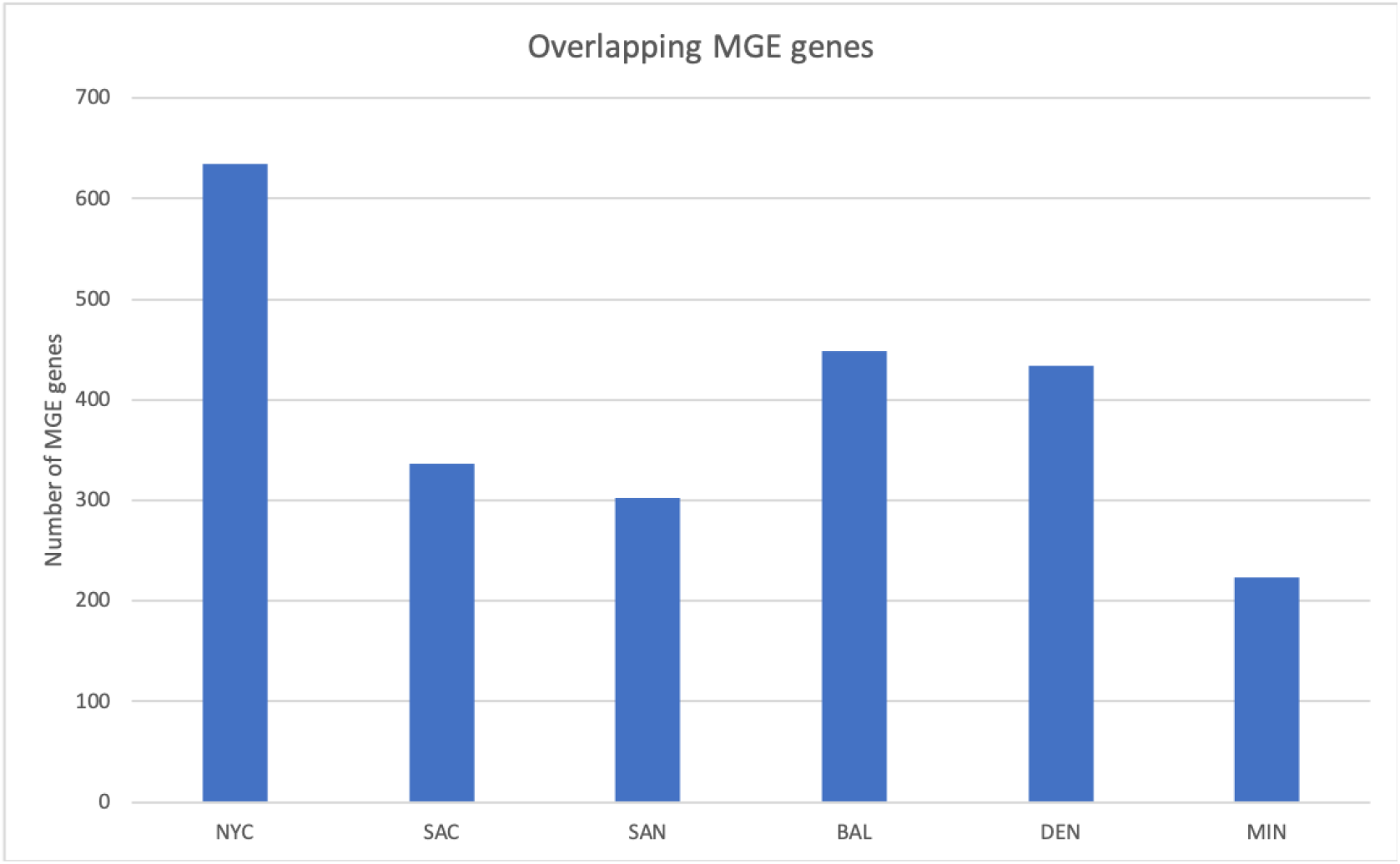
The bar plot shows the distributions of overlapping MGE genes among the US cities.

#### 3.1.2 On the virome

A total of 70,839 viral contigs were detected; however, only 394 contigs (0.56%) were considered relevant by CheckV (Medium-quality, High-quality, and Complete). Upon investigation, it was observed that the number of viral contigs was affected by the small-sized contigs generated by the assembly. VirFinder was responsible for over 95% of the total number of viral contigs detected, with a high false positive rate. We inferred the high false positive rate based on the quality control performed by the CheckV tool. Putative viral contigs detected by VirFinder that lacked viral genes were tagged as non-viral and counted as falsely identified as viruses.

#### 3.1.3 On the mobilome

Table 4 shows the distribution of MGE genes across all samples. We identified a total of 1660 MGE genes across 6 US cities, and Figure 4 shows the number of same MGE genes observed across different US cities. For example, a total of 142 genes were found to be present in all US cities, whereas 501 genes were only found once in each city with varying frequency. This indicates that there are some MGE genes common among across cities, while some are city-specific.

**Figure 4.**
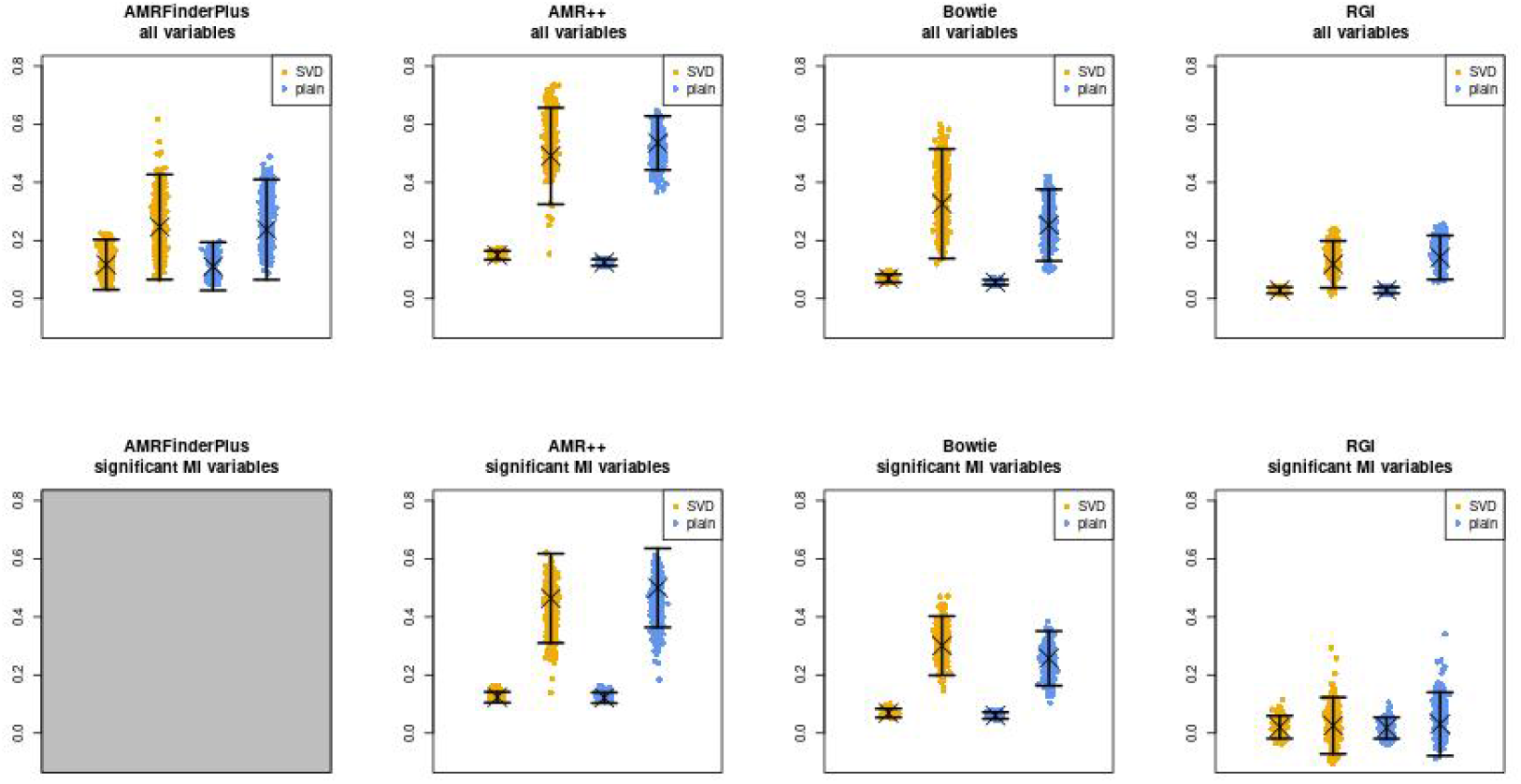
Estimates of *S_in_ − S_btw_*, for datasets generated by different tools (columns) and similarity variants (colors). Y axis of each plot is the value of the statistic. Each X position corresponds to each variant. Left of each color corresponds to statistic computed on whole similarity matrix. Right corresponds to statistic computed on sparsified matrix. ’X’ marks the value computed on real data, jitter plots show distribution of the jackknife replicates. Error bars signify standard errors. Top row shows results of calculation using all markers, bottom one – limited to markers significant to ’city’ variable by MI test.

**Table 3.**
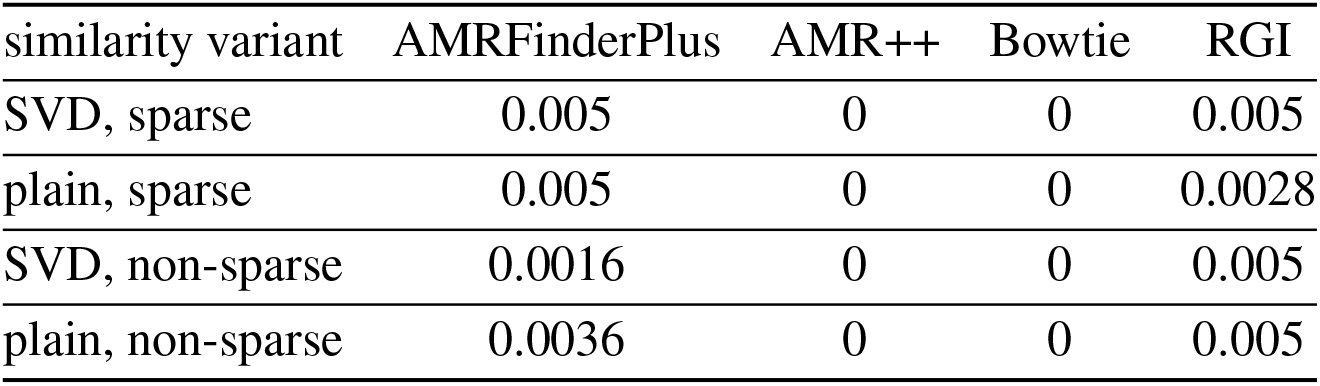
Holm corrected p-values of the permutation test on the *S_in_ − S_btw_* statistic. All similarity variants were calculated using all markers (without filtering relevant ones).

**Table 4.** Distribution of Mobile Genetic Elements (MGE) Genes Across Cities. This table shows the number of samples, genes, and protein hits (homologs) for each city analyzed in the study.

| City | samples | genes | protein_hits (homologs) |
| --- | --- | --- | --- |
| DEN | 45 | 924 | 8279 |
| BAL | 14 | 910 | 8184 |
| SAC | 16 | 867 | 7248 |
| NYC | 46 | 1577 | 32798 |
| SAN | 16 | 666 | 5051 |
| MIN | 6 | 532 | 3943 |

Among the MGE genes we identified, 438 MGE’s genes co-occur with 325 AMR markers across the US cities, and a total of 12 of the MGE-AMR gene marker patterns were found across all cities (Table 5). A detailed table about the co-occurrence between the MGE-AMR across US cities with city and sample information can be found in the supplementary materials. The Table 5 shows most common MGE gene and AMR gene that co-occured together among all 6 cities of the US.

**Table 5.** Prevalence of Mobile Genetic Elements (MGE) and Antibiotic Resistance Markers (AMR) Across US Cities. This table lists the most common MGE and AMR genes, along with the number of cities, samples, and sequencing reads in which they were identified. Verification ensures that genes listed are distinct despite similar names.

| MGE | AMR | Cities | Samples | Fastq reads |
| --- | --- | --- | --- | --- |
| gyrA | GYRA | 6 | 116 | 3819 |
| parC | PARC | 6 | 66 | 1271 |
| parE | PARE | 6 | 93 | 2628 |
| gyrB | GYRB | 6 | 73 | 3601 |
| dnaK | DNAK | 6 | 75 | 288 |
| gyrB | PARY | 6 | 33 | 75 |
| tnpA_IS6100 | VEB | 6 | 9 | 19 |
| tnpA | VEB | 6 | 41 | 151 |
| gyrB | GYRBA | 6 | 69 | 1559 |
| thyA | THYA | 6 | 20 | 44 |
| ruvB | RUVBM | 6 | 18 | 37 |
| gyrB | GYRBZ | 6 | 7 | 7 |

### 3.2 Clustering

Figure 4 shows the point estimates of the similarity comparison statistic. Table 3 contains the FWER corrected p-values for the *S_in_ − S_btw_* statistic. Overall, results suggest that based on AMR levels obtained from both AMR++ and Bowtie derived datasets we can find that in fact similarities of samples coming from the same cities are greater than similarities of samples coming from different cities. Comparing the point estimates for those tools (first row of fig. 4) shows that in terms of finding markers relevant for geographic location, AMR++ performs slightly better. Significant difference for the sparsified variants also suggests that indeed, the strongest similarities form along samples from the same cities, at least for data obtained with AMR++ and Bowtie methods.

AMRFinderPlus and RGI in that test performed worse, which is both visible in p-values of the permutation test as well as on the point estimate plots. For those tools, the difference between sparsified and normal similarity variant is within the bounds of standard error, which reinforces that observation. ’RGI’ came up as the worst from all of the used tools.

Limitation to the significant MI variables does not seem to bring much difference. In the end, AMR++ stands out as the best approach, without any significant difference between SVD based similarity and plain one.

We show the visualization of the similarities for the SVD embedded samples derived from the AMR++ dataset (fig. 5). We see that samples from different cities still have strong connections to each other, but blocks along the diagonal are noticeable. Overall, unsupervised analysis highlighted the need for including information external to the clustering procedure.

**Figure 5.**
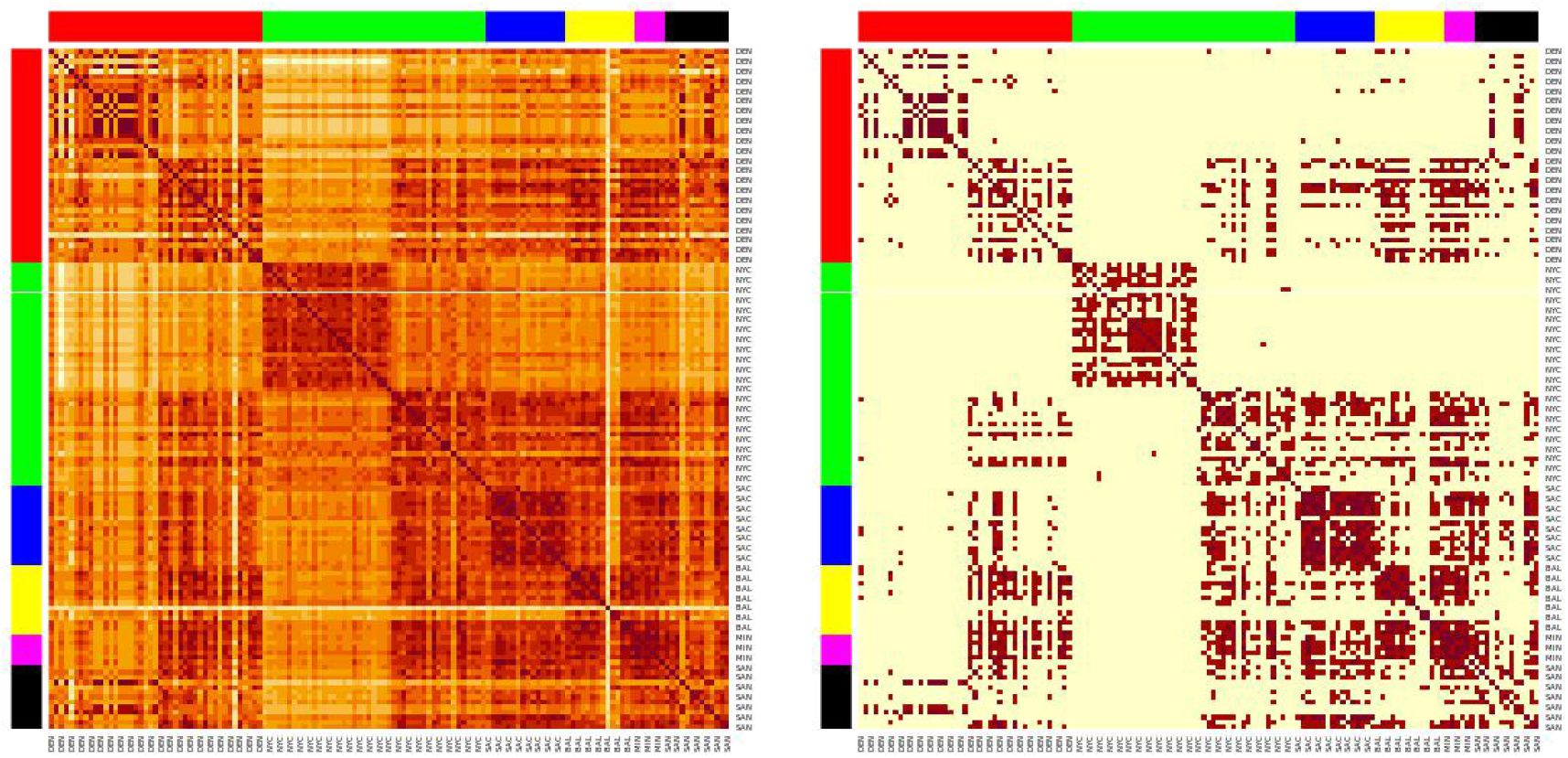
Heatmaps of the sample similarities for the AMR++ derived dataset, using the SVD embedding applied on significant MI markers. Right shows sparsified matrix, left – the plain one. Each row/column corresponds to each sample. Row/columns are arranged by the city labels, which are also marked by the side colors. Cities are arranged in following order (left to right): DEN, NYC, SAC, BAL, MIN, SAN

SAC samples (blue colors) form a noticable cluster, as seen on both sparsified and plain version of the heatmap plot. This corresponds to the high classification score by ML model for that particular city, as we show in the next section. We can also spot a well separated cluster in the middle that corresponds to portion of samples coming from NYC (green colors). Baltimore group (yellow) contains several outliers visibly distinct from the rest. Both of these features turned out to be partly explained by k-mer content of each sample as which we show later.

### 3.3 **K-mer Profiling Results**

K-mer profiling revealed significant insights into the diversity and complexity of the metagenomic samples. As shown in Figure 6, each row and column corresponds to a sample. The line plots on top of the figure represent the number of unique k-mers at each X coordinate (sample). The yellow, orange, and red curves represent k-mer sizes 33, 55, and 77, respectively. The colors in the margins represent samples from different cities: DEN (red), NYC (green), SAC (blue), BAL (yellow), MIN (magenta), and SAN (black).

**Figure 6.**
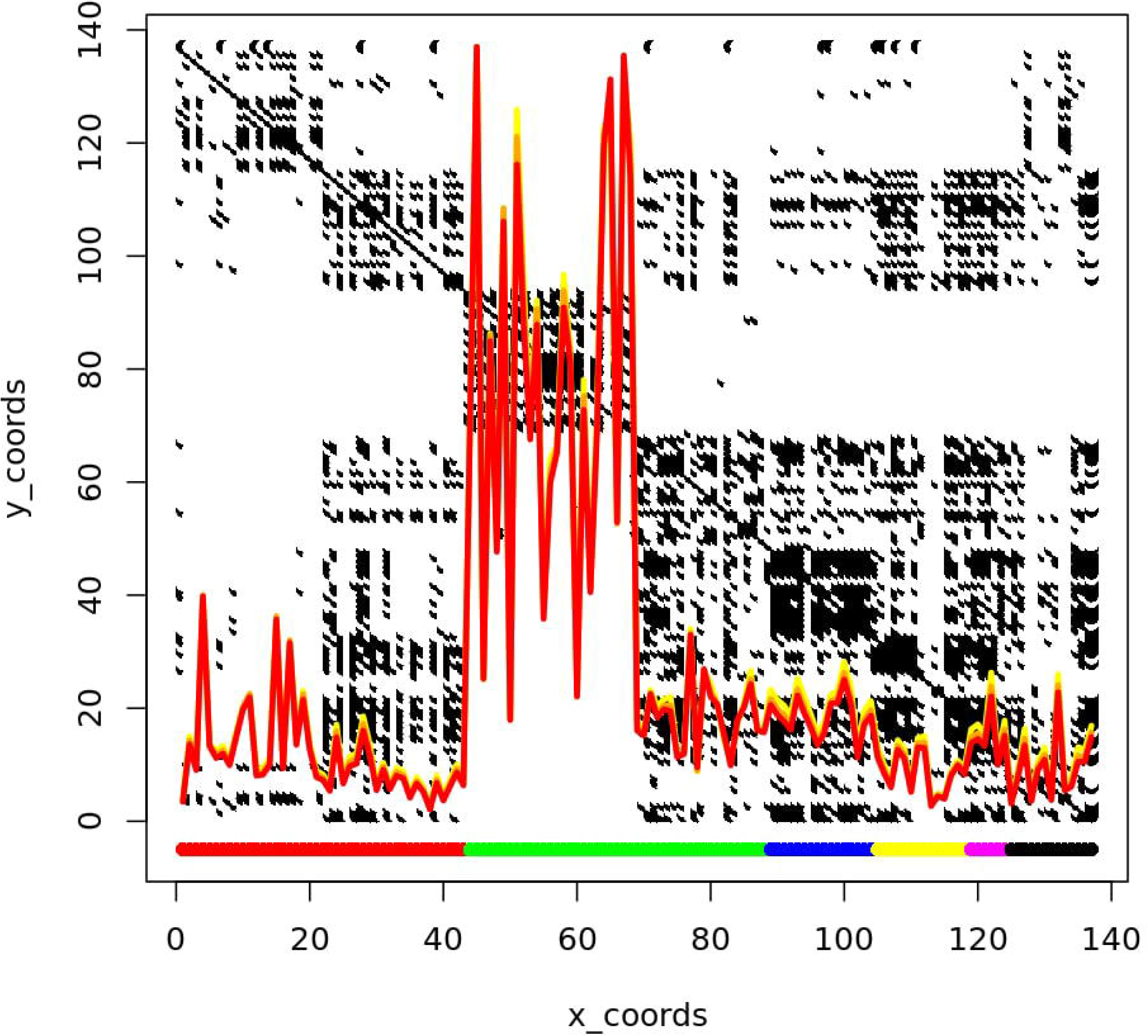
The line plots represent the number of unique k-mers at each X coordinate, with the yellow, orange, and red curves corresponding to k-mer sizes 33, 55, and 77, respectively. The margin colors indicate samples from different cities: DEN (red), NYC (green), SAC (blue), BAL (yellow), MIN (magenta), and SAN (black). The peak k-mer counts are notably higher for New York City (NYC) samples, which are divided into two clusters: one with high complexity containing only NYC samples, and one with low complexity, shared with samples from Sacramento (SAC) and potentially Baltimore (BAL).

**Figure 7.**
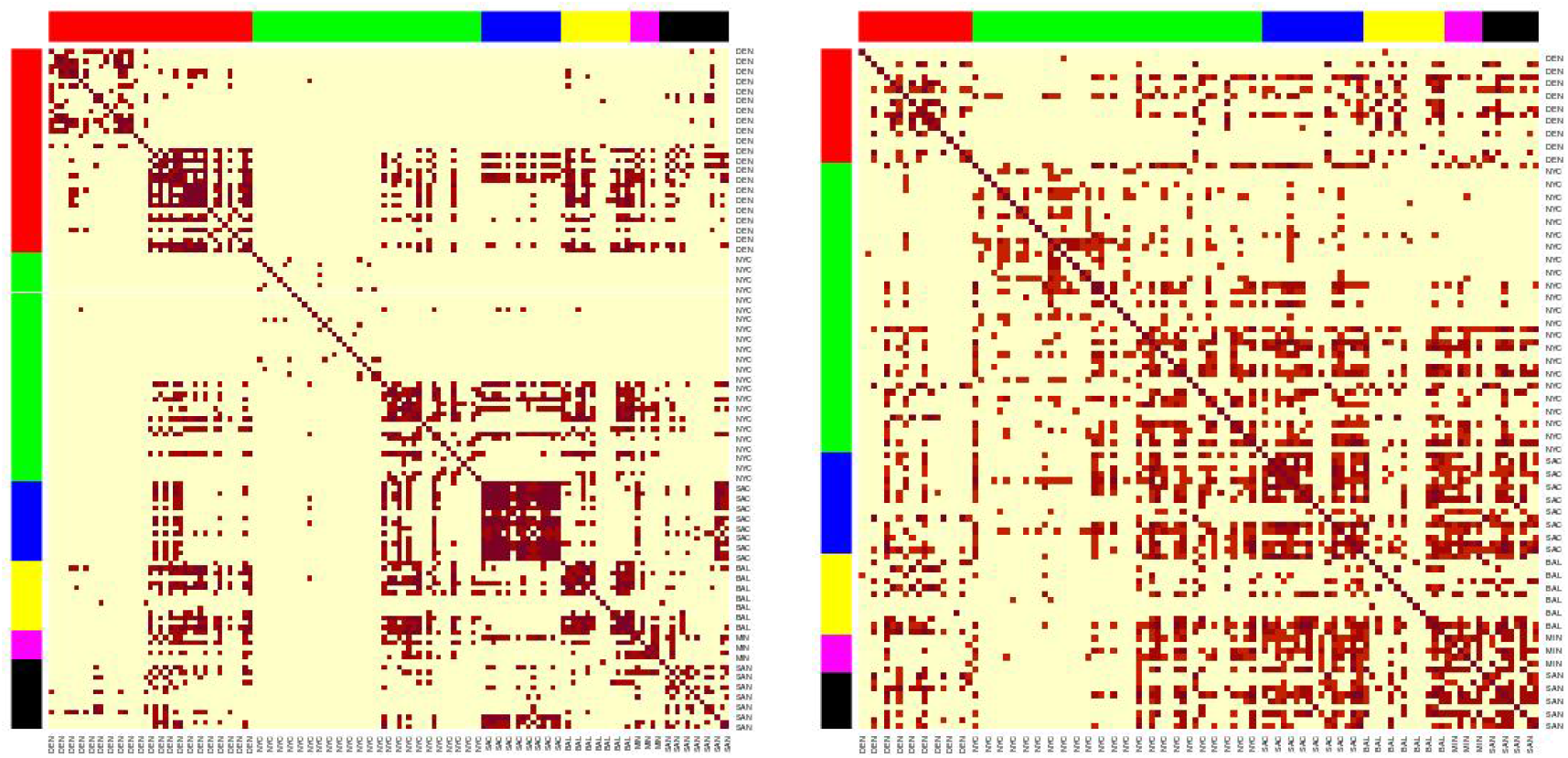
Heatmaps of the sample similarities for the AMR++ derived dataset, using the SVD embedding applied on significant MI markers. Left(50% MGE filtering) shows clear clusters, while right(100% MGE filtering) no longer shows clusters after removing mAMRs. Each row/column corresponds to each sample. Row/columns are arranged by the city labels, which are also marked by the side colors. Cities are arranged in following order (left to right): DEN, NYC, SAC, BAL, MIN, SAN

As observed, the peak of k-mer counts is particularly pronounced for samples from New York City (NYC). NYC samples are split into two distinct clusters, one with high complexity and one with low complexity. The high complexity cluster is exclusively composed of NYC samples, while the low complexity cluster includes samples from Sacramento (SAC) and potentially Baltimore (BAL). This clustering pattern suggests that elevated k-mer counts from NYC samples may be due to shared genetic elements or contamination, leading to unexpected clustering across different cities.

### 3.4 **Modelling**

The four tools and the six normalization approaches were tested. The multi-class models yielded poor results. The likely reason was the large number of classes with a small number of in-class samples.

Therefore, we have focused on pair classification. Those were done in one-vs-all and one-vs-one manner and can be seen in table 7. Notably, Bowtie and AMR++ displayed the best results, indicating that reads-based resistome profiling is more accurate than assembled-based profiling. Bowtie and AMR++ consistently demonstrated the highest performance across the pairwise classifications. Specifically, in the one-vs-all evaluations, Bowtie and AMR++ achieved average AUC scores of 0.94. Similarly, in the one-vs-one comparisons, Bowtie and AMR++ maintained high performance, with average AUC scores of 0.92 and 0.94, respectively. This indicates that reads-based resistome profiling, utilized by Bowtie and AMR++, tends to be more accurate than assembly-based approaches, as evidenced by their superior AUC values. The results reveal a significant deviation from randomness (an AUC of 0.5), demonstrating the effectiveness of Bowtie and AMR++ in accurately profiling the resistome.

**Table 6.** Prediction Accuracy for AMR++ with MGE Removal Scenarios. This table compares the accuracy when Mobile Antimicrobial Resistance Markers (mAMRs) are removed under two conditions: ‘Removal MGE 50cov’ where mAMRs in at least 50% present, and ‘Removal all MGE’ where all mAMRs are excluded. Results are shown for one-versus-one and one-versus-all city comparisons to assess the impact of MGE on predictive accuracy.

|  | removal MGE 50cov | Standard | removall all MGE |
| --- | --- | --- | --- |
| **one versus one** |  |  |  |
| DEN BAL | 0.9247 | 0.9869 | 0.9231 |
| MIN BAL | 1.0000 | 0.9556 | 0.9487 |
| MIN DEN | 1.0000 | 0.9941 | 0.9259 |
| NYC BAL | 0.8947 | 0.9327 | 0.8979 |
| NYC DEN | 1.0000 | 1.0000 | 0.9221 |
| NYC MIN | 0.9211 | 0.9636 | 0.9316 |
| SAC BAL | 1.0000 | 1.0000 | 0.7835 |
| SAC DEN | 0.9924 | 0.9958 | 0.8750 |
| SAC MIN | 1.0000 | 0.9714 | 1.0000 |
| SAC NYC | 1.0000 | 1.0000 | 0.8329 |
| SAN BAL | 0.8951 | 0.9877 | 0.9402 |
| SAN DEN | 0.8615 | 1.0000 | 0.9568 |
| SAN MIN | 0.9692 | 0.9556 | 0.9815 |
| SAN NYC | 0.9798 | 1.0000 | 0.8913 |
| SAN SAC | 0.9256 | 1.0000 | 0.9583 |
| Average | 0.9577 | 0.9861 | 0.9181 |
| **one versus all** |  |  |  |
| BAL versus all | 0.8709 | 0.9520 | 0.5887 |
| DEN versus all | 0.9359 | 0.9088 | 0.8129 |
| MIN versus all | 0.9571 | 0.9890 | 0.8660 |
| NYC versus all | 0.9611 | 0.9748 | 0.8683 |
| SAC versus all | 0.9831 | 0.9944 | 0.7378 |
| SAN versus all | 0.8957 | 0.9474 | 0.6891 |
| Average | 0.9340 | 0.9611 | 0.7605 |

**Table 7.** Comparison of Prediction Results by AMR Detection Tools. This table presents the accuracy scores for AMRFinder, RGI, Bowtie, and AMR++ in one-versus-all and one-versus-one prediction scenarios across different cities.

|  | AMRFinder | RGI | Bowtie | AMR++ |
| --- | --- | --- | --- | --- |
| **one versus all** |  |  |  |  |
| BAL versus all | 0.69 | 0.71 | 0.94 | 0.93 |
| DEN versus all | 0.81 | 0.89 | 0.92 | 0.91 |
| MIN versus all | 0.64 | 0.89 | 0.89 | 0.97 |
| NYC versus all | 0.96 | 0.92 | 0.97 | 0.96 |
| SAC versus all | 0.60 | 0.78 | 0.94 | 0.99 |
| SAN versus all | 1.00 | 0.34 | 0.96 | 0.86 |
| Average | 0.78 | 0.75 | 0.94 | 0.94 |
| **one versus one** |  |  |  |  |
| DEN BAL | 0.66 | 0.74 | 0.96 | 0.92 |
| MIN BAL | 0.96 | 0.96 | 0.82 | 0.99 |
| MIN DEN | 0.79 | 0.98 | 0.89 | 1.00 |
| NYC BAL | 0.69 | 0.92 | 0.97 | 0.92 |
| NYC DEN | 0.98 | 0.98 | 0.98 | 0.99 |
| NYC MIN | 1.00 | 0.98 | 0.93 | 0.96 |
| SAC BAL | 0.99 | 0.83 | 0.99 | 1.00 |
| SAC DEN | 0.98 | 0.93 | 0.94 | 0.99 |
| SAC MIN | 1.00 | 0.95 | 0.99 | 0.99 |
| SAC NYC | 0.91 | 0.88 | 0.99 | 0.99 |
| SAN BAL | 0.00 | 0.76 | 0.97 | 0.93 |
| SAN DEN | 0.00 | 1.00 | 0.86 | 0.88 |
| SAN MIN | 0.00 | 0.91 | 0.80 | 0.98 |
| SAN NYC | 0.00 | 0.92 | 0.99 | 0.99 |
| SAN SAC | 0.00 | 0.87 | 0.91 | 0.93 |
| Average | 0.70 | 0.89 | 0.92 | 0.94 |

The resistome profile generated by AMR++ was used for further experiments. The reason was manyfold: 1) It had the highest and most diverse number of AMRs detected, 2) the highest number of AMRs detected that matched the AMRs present in the isolates, 3) the highest inter-city dissimilarity, and 4) higher AUC values for the different modelling approaches. Table 8 displays the AUC values for different normalization approaches using AMR++ data. The values found on upper part of the table display how distinguishable is a given city from the rest of the dataset while the values on the lower part display how distinguishable is a city from another, pairwise. Each column represents a different pre-processing approach. The ”Standard” column shows the values directly outputted by AMR++. The ”binary” column represents the binary version of these values, where the data is converted into binary form, indicating the presence or absence of features. The best-preprocessing approach was the binarization. It has show the highest values across all comparisons, followed by Standard ouput. Normalization by the number of SSU units (SSU total and SSU cov50) did not significantly improve modelling results. The same was observed for normalization based on sequencing and assembly nucleotides.

**Table 8.** Comparative Prediction Accuracy of AMR++ Tool. This table presents the prediction accuracy results of the AMR++ tool across various normalization modes, including standard, binary, SSU total, SSU cov50, reads, and genome data. The data is shown for both one-versus-all and one-versus-one city comparison scenarios.

|  | standard | binary | SSU total | SSU cov50 | reads | Genome |
| --- | --- | --- | --- | --- | --- | --- |
| **one versus all** |  |  |  |  |  |  |
| DEN vs all | 0.92 | 0.93 | 0.93 | 0.94 | 0.93 | 0.94 |
| NYC vs all | 0.92 | 0.97 | 0.95 | 0.96 | 0.95 | 0.94 |
| SAC vs all | 0.91 | 0.95 | 0.93 | 0.94 | 0.95 | 0.94 |
| BAL vs all | 0.86 | 0.86 | 0.91 | 0.89 | 0.89 | 0.89 |
| MIN vs all | 0.93 | 0.98 | 0.96 | 0.99 | 0.96 | 0.96 |
| SAN vs all | 0.86 | 0.81 | 0.91 | 0.85 | 0.85 | 0.85 |
| **one versus one** |  |  |  |  |  |  |
| DEN - NYC | 0.99 | 0.96 | 0.94 | 0.97 | 0.95 | 0.96 |
| DEN - SAC | 0.96 | 0.97 | 0.93 | 0.95 | 0.94 | 0.95 |
| DEN - BAL | 0.96 | 0.98 | 0.97 | 0.95 | 0.97 | 0.97 |
| DEN - MIN | 0.94 | 0.97 | 0.95 | 0.95 | 0.94 | 0.95 |
| DEN - SAN | 0.87 | 0.86 | 0.88 | 0.90 | 0.89 | 0.90 |
| NYC - SAC | 0.94 | 0.98 | 0.96 | 0.97 | 0.96 | 0.96 |
| NYC - BAL | 0.97 | 0.99 | 0.99 | 0.98 | 0.98 | 0.98 |
| NYC - MIN | 0.99 | 0.99 | 0.97 | 0.98 | 0.98 | 0.98 |
| NYC - SAN | 0.95 | 0.98 | 0.96 | 0.97 | 0.96 | 0.97 |
| SAC - BAL | 0.97 | 1.00 | 0.98 | 0.99 | 0.99 | 0.99 |
| SAC - MIN | 0.96 | 1.00 | 0.97 | 0.99 | 0.99 | 0.99 |
| SAC - SAN | 0.90 | 0.93 | 0.91 | 0.92 | 0.92 | 0.92 |
| BAL - MIN | 0.93 | 0.93 | 0.90 | 0.92 | 0.91 | 0.92 |
| BAL - SAN | 0.90 | 0.93 | 0.91 | 0.92 | 0.91 | 0.92 |
| MIN - SAN | 0.98 | 1.00 | 0.98 | 1.00 | 1.00 | 1.00 |

### 3.5 **Linking AMRs to MGEs and viruses**

Aiming to observe how mobile genetic elements (MGEs) and viruses influence antimicrobial resistance, we excluded antimicrobial resistance genes (AMRs) associated with these elements and repeated the classification process. This involved removing virus-associated AMRs (vAMRs) and mobile genetic element-associated AMRs (mAMRs) through a two-step process. First, we used Bowtie2 mapping to identify reads linked to either viruses or MGEs. Then, we examined the remaining reads using the AMR++ tool. Out of nearly 70,000 viral contigs detected, only 4 were associated with resistance markers, all belonging to New York City. Therefore, we decided to utilize only mAMRs in the downstream analysis.

### 3.6 **Filtering mAMRs**

We analyzed MGEs observed in more than one city, focusing on their proportions in each location. When selecting MGEs to filter, we considered two criteria: 1) MGEs must be observed in more than one city, and 2) each city must have a minimum presence of 30%, 50%, 80%, or 100%. Additionally, we selected MGEs for removal by randomly removing reads equal to the number removed during the 50% filtering criteria from each sample. These filtering criteria were used to identify potential candidates for mAMRs. The mAMRs were then filtered at the FASTQ read level by removing reads containing the selected MGEs. Subsequently, we recalculated the antibiotic resistance markers (AMRs) using the AMR++ tool.

We utilized the newly generated antibiotic resistance matrix from the AMR++ detection to reassess AMR city similarity analysis and modeling. Among the four MGE filtering percentages, the 30% criterion resulted in minimal changes, while the 100% criterion led to a complete reshuffling of antibiotic resistance (AMR) clusters and deterioration in model performance. The 50% criterion showed some improvements in the models, although the AMR clusters exhibited minimal variation, as our analysis was based on binarized information. Random removal of MGEs did not affect the results of AMR city similarity analysis and models.

These results suggest that certain resistome markers are inherently connected with mobile genetic elements. The removal of these markers significantly alters the resistome profile, impacting the efficiency of resistome modeling. Therefore, careful consideration must be given when filtering MGEs, as their presence or absence can drastically change the resistome profile and affect the accuracy of AMR detection and modeling. Our findings highlight the importance of preserving key MGEs in resistome studies to maintain the integrity and predictive power of resistome profiling models.

## 4 CLOSURE

### 4.1 **Conclusion and Discussion**

AMR exhibits a heterogeneous distribution across the dataset, with varying resistome profiles that do not correlate with sample depth. The investigated samples did not present nearly half of the ARGs presented in the isolates, indicating that either: 1) the sequencing depth of the urban samples was insufficient, 2) the isolated species were not dominant in the urban dataset, or 3) the classification methods were limited by incomplete reference databases.

Statistical analysis of the similarities suggested that the link between the origin of the sample and its AMR levels is non-random. In regards to how much of location-relevant markers each tool finds, we can see non-negligible variability between approaches, highlighting AMR++ and Bowtie as most informative in that aspect. Regardless of all of these general considerations, in practice we still see pairs of samples coming from different cities which are more strongly connected than pairs from the same city, which means that inferring their origin based on the clustering would not be effective. For some of the cities, such as DEN and SAC we can see that all of the samples are tightly connected. However, for clusters cross city boundaries and inside some of the cities we can spot subgroups not related to each other at all (e.g., DEN and NYC). This fact seems to correspond to different counts of total k-mer content, especially evident by split within NYC group. Interestingly, removal of MGE related reads leads to disappearance of the clusters, which highlights their importance to the study of AMRs.

#### 4.1.1 Modelling

AMR++ has proven to be the best tool for resistome modeling. It relies on quality-controlled reads rather than assemblies, allowing it to detect the highest and most diverse number of AMR markers. Additionally, it achieves the highest AUC scores in random forest prediction of sampling sites. This is encouraging news for newcomers in the field, as the assembly process can be a significant bottleneck due to its high demand for computational resources.

Binarization appears to be the most effective data pre-processing approach because it simplifies the data, reduces noise and overfitting, and enhances the detection of critical signals by focusing on the presence or absence of resistance genes rather than their abundance. This method generally achieved higher AUC scores, suggesting improved classification performance and predictive power

#### 4.1.2 Isolates prediction

Classification proved to be challenging, with multiclass prediction resulting in many misclassifications. AMR++ was the only tool that produced noteworthy results, but they were still incorrect. Limitations of the database and the uneven number of samples for each city skewed predictions towards NYC. Efforts to clean the dataset by removing AMR markers associated with viruses or MGEs influenced the modeling but did not improve the predictions.

### 4.2 **Limitations and future work**

While this study has provided valuable insights into the distribution and diversity of AMRs in urban microbiomes, future studies could achieve more robust results by including a larger sample size, deeper sequencing, and a wider range of AMR identification tools and databases, along with additional metadata. These efforts could offer a more comprehensive understanding of the AMR landscape in urban environments and the effects of anthropogenic influence, thereby fostering the development of effective prevention strategies.

## AUTHOR CONTRIBUTIONS

R.B.T. was responsible for experiment design, data preprocessing, data generation, data analysis, and drawing conclusions. W.L. focused on conducting random forest experiments and subsequent prediction analyses. P.S. handled clustering analyses. B.S. contributed to data generation and data analysis. W.R.R. and P.Ł. provided overall coordination and supervision of the project.

## ACKNOWLEDGMENTS

We gratefully acknowledge Poland’s high-performance Infrastructure PLGrid - ACK Cyfronet AGH for providing computer facilities and support within computational grant no PLG/2023/016299. Special thanks also go to the high-performance cluster at the Małopolska Centre of Biotechnology of Jagiellonian University, and to its dedicated system administrator, Wojciech Pilch, for his invaluable support in our computational endeavors.

## DATA AVAILABILITY STATEMENT

The datasets analyzed for this study can be found in the CAMDA 2023 Challenge (2023).CAMDA 2023 challenge page: http://camda2023.bioinf.jku.at/contest dataset. All scripts and pre-processed tables with results can be found a dedicated Github repository: https://github.com/rbtoscan/frontiers camda 2023.

