## supplementary file for "Antimicrobial Resistance in Diverse Urban Microbiomes: Uncovering Patterns and Predictive Markers"

### Supplementary Material

#### 1 CLUSTERING PROCEDURE DETAILS

##### 1.1 Sparisifcation step in detail

After computing the cosine similarity, in part of the variants, weaker connections were zeroed out using the threshold chosen according to the criterion of maximum number of cliques of size at least 3 found on the thresholded similarity matrix. This is a heuristic for choosing the sparsification threshold dependent on the network's topology characteristics induced by each specific similarity matrix, which we found empirically to work well in various scenarios. For a dataset of $N$ samples, we get a $N \times N$ similarity matrix $S$ of non-negative entries, where $S_{ij}$ encodes the degree of the similarity between objects $i, j$. Taking $S$ as a weight matrix of the weighted graph, we find its partition into heavily connected disjoint cliques by an efficient greedy algorithm and count them. We count such cliques on a thresholded $S^{thr}$ version of the similarity matrix $S$. Finally, we pick the final sparsification threshold $thr$, which maximizes the number of cliques of size at least 3, from all of the tested candidate thresholds.

##### 1.2 Statistics of the permutation test

Null distribution in a permutation test is a distribution over all possible rearrangements of city labels. Exact p-value of a statistic $\widehat{\theta_{real}}$ calculated on a real dataset can then be calculated as (for one-sided test) Efron and Tibshirani (1993):

$$\frac{\#\text{rearrangements with } \widehat{\theta_{rearrangement}} \geq \widehat{\theta_{real}}}{\#\text{all rearrangements}} \tag{S1}$$

If specific value of $\widehat{\theta_{real}}$ is not special, then any other rearrangement would be likely to produce a statistic of the size like $\widehat{\theta_{real}}$ or bigger (in the case when big value of $\widehat{\theta_{real}}$ is meaningful as in the case with the AMR based inside vs between city similarity difference). Then the number of rearrangements with a value of the statistic exceeding the $\widehat{\theta_{real}}$ would be big.

If the $\widehat{\theta_{real}}$ estimate we have from the data is non-random, such that the difference between AMR based similarities truly comes from geographical differences, then number of such rearrangements would be small. Then such a p-value is also small.

##### 1.3 Computational details of the clustering protocol

For selecting relevant variables in the 'data limitation before cosine calculation' step, we have used a threshold of significance of 0.05 with a Benjamini–Yekutieli correction.

For SVD embedding dimension, we have arbitrarily chosen the $\frac{1}{3}$ of the maximum possible dimension for each dataset. It has to be noted here that cosine similarity between two vectors $v, w$ is meaningful only if both of those vectors have norm $> 0$. Therefore, before computing the cosine similarity itself, one has to filter out samples which have zero norm. In some of the variants, transformations can induce the zero norm on some of the original samples themselves. For that reason, number of used samples for each dataset and similarity variant can differ slightly. In case of the tool 'AMRFinderPlus' and limitation of variables to the subset significantly related to the 'city' variable, the number of non-zero norm samples was reduced to 52 (vs 92 originally available). Since these variants turned out to be difficult to compare with the rest, we do not report statistics for them.

For the jackknife standard error estimation, to compute each replicate we have left out $0.25$ of the original dataset (in a stratified manner described before). Choice of such fraction follows both from the theory (Shao and Wu (1989)) (such that number of left out samples, $d$, is contained in $(\sqrt(N), N)$) and class distribution of the 'city' variable (smallest size of the smallest class across all variants is 4). Number of replicates for the jackknife estimation was equal to 500.

Lastly, for the permutation test, we have used 5000 rearrangements and we have limited ourselves to the variants where we use all of the available markers for cosine calculation. Reason for the latter is that when we permute the labels of the 'city' variable, it is very unlikely to find any marker relevant for it. For these rearrangements one cannot compute the similarity and this makes the comparisons between variants difficult to execute fairly.

#### 2 SUPPLEMENTARY DATA

Supplementary Material should be uploaded separately on submission. Please include any supplementary data, figures and/or tables. All supplementary files are deposited to FigShare for permanent storage and receive a DOI.

Supplementary material is not typeset so please ensure that all information is clearly presented, the appropriate caption is included in the file and not in the manuscript, and that the style conforms to the rest of the article. To avoid discrepancies between the published article and the supplementary material, please do not add the title, author list, affiliations or correspondence in the supplementary files.

#### 3 SUPPLEMENTARY TABLES AND FIGURES

For more information on Supplementary Material and for details on the different file types accepted, please see the Supplementary Material section of the Author Guidelines.

Figures, tables, and images will be published under a Creative Commons CC-BY licence and permission must be obtained for use of copyrighted material from other sources (including re-published/adapted/modified/partial figures and images from the internet). It is the responsibility of the authors to acquire the licenses, to follow any citation instructions requested by third-party rights holders, and cover any supplementary charges.

##### 3.1 Figures
